# Interdependent Phenotypic and Biogeographic Evolution Driven by Biotic Interactions

**DOI:** 10.1101/560912

**Authors:** Ignacio Quintero, Michael J. Landis

## Abstract

Biotic interactions are hypothesized to be one of the main processes shaping trait and biogeographic evolution during lineage diversification. Theoretical and empirical evidence suggests that species with similar ecological requirements either spatially exclude each other, by preventing the colonization of competitors or by driving coexisting populations to extinction, or show niche divergence when in sympatry. However, the extent and generality of the effect of interspecific competition in trait and biogeographic evolution has been limited by a dearth of appropriate process-generating models to directly test the effect of biotic interactions. Here, we formulate a phylogenetic parametric model that allows interdependence between trait and biogeographic evolution, thus enabling a direct test of central hypotheses on how biotic interactions shape these evolutionary processes. We adopt a Bayesian data augmentation approach to estimate the joint posterior distribution of trait histories, range histories, and co-evolutionary process parameters under this analytically intractable model. Through simulations, we show that our model is capable of distinguishing alternative scenarios of biotic interactions. We apply our model to the radiation of Darwin’s finches—a classic example of adaptive divergence—and find support for *in situ* trait divergence in beak size, convergence in traits such as beak shape and tarsus length, and strong competitive exclusion throughout their evolutionary history. Our modeling framework opens new possibilities for testing more complex hypotheses about the processes underlying lineage diversification. More generally, it provides a robust probabilistic methodology to model correlated evolution of continuous and discrete characters.

---

One of the major goals of biogeography is to explain the dramatic variation in species richness across the planet. Ultimately, any difference in species richness between two regions stems from contrasting frequencies of speciation, extinction or dispersal events (Ricklefs 1987). While diversification processes alone drive the total number of species through time, range evolution dynamics cannot be ignored when explaining spatial gradients of biodiversity (Wiens and Donoghue 2004). Indeed, the increase in richness within an area can only be the result of a new species eventually coming into (or remaining in) sympatry (Weir and Price 2011; Pigot and Tobias 2013). This necessarily involves two general processes: that of lineage splitting followed by that of establishing coexistence. Yet, we still lack a basic understanding on the generality and magnitude of the different processes that shape the geographical and phenotypic evolution of diversifying lineages (Mayr 1970; MacColl 2011; Tobias et al. 2014; Clarke et al. 2017).

Evidence suggests that the great majority of speciation processes, at least in terrestrial animals, involve an allopatric phase, with few conclusive examples demonstrating parapatric or sympatric speciation in nature (Mayr 1970; Coyne and Orr 2004; Rundell and Price 2009), but see (Stroud and Losos 2016). The prevailing view asserts that new species arise from geographically isolated populations that evolve sufficient morphological, ecological, physiological, behavioral and/or genetic differences to act as reproductive barriers. These incipient species usually fill very similar ecological niches since the initial driver of reproductive isolation was chance separation by geographical barriers (Kozak and Wiens 2006; Rundell and Price 2009; Cadena et al. 2011; Smith et al. 2014). Equivalent ecological requirements are supposed to make long-term coexistence untenable, following the competitive exclusion principle (Gause 1934; Hardin 1960; Macarthur and Levins 1967). Recent radiations often follow this principle, with closely related species occupying similar habitats but separated by physical barriers (recognized more than one century ago as the “general law of distribution”; Jordan 1905; Rundell and Price 2009). For species to attain sympatry, and thus elevate local richness, coexistence theory predicts that species must diverge sufficiently along one or more niche axes to avoid competition (Elton 1946; Hardin 1960; Macarthur and Levins 1967; Diamond 1978; Grether et al. 2009; Godoy et al. 2014).

Consequently, biotic interactions seem to be paramount in shaping trait and biogeographic distributions of evolving lineages. The effects of biotic interactions during evolutionary radiations can be broadly categorized in three ways: by limiting (or enhancing) geographical expansion (Rundell and Price 2009; Ricklefs 2010; Weir and Price 2011; Pigot and Tobias 2013; Tobias et al. 2014; Pigot et al. 2018), by promoting (or reducing) local extinction (Slatkin 1974; Simberloff and Boecklen 1991; Valone and Brown 1995), and by inducing niche divergence (or convergence) in coexisting species (Lack 1954; Rohwer 1973; Schluter 2000; Davies et al. 2007; Pfennig and Pfennig 2012). While there are experimental tests and suitable models for shallow divergences under population genetic or ecological models (e.g., Lotka 1924; Neuhauser and Pacala 1999; Schluter 2000; Scheffer and van Nes 2006), the long-term evolutionary consequences of biotic interactions measured at ecological time-scales remain difficult to characterize. Except for a few illuminating—but serendipitous—fossil sequences (Elredge 1974; Schindel and Gould 1977), our understanding has been mostly restricted to tests of phylogenetic community structure metrics, such as measures of trait under/over-dispersion juxtaposed to null models (Webb et al. 2002; Cavender-Bares et al. 2009), and correlative analyses, such as sister-species comparisons between allopatric species and those that have achieved secondary sympatry (Schluter et al. 1985; Davies et al. 2007; Pigot and Tobias 2013; Anacker and Strauss 2014; Freeman 2015; Cadotte et al. 2017; McEntee et al. 2018). Though insightful, such pattern-based studies rely on non-generative models that do not disentangle how the processes are driven by biotic interactions over evolutionary timescales. The different stages of biotic interactions unfold in a complex interplay between phenotype and geographical distribution, often ephemeral through the evolutionary history of species (Brown and Wilson 1956), and most probably lost when evidence is restricted to contemporaneous observations (Schindel and Gould 1977). To understand this interplay, generative phylogenetic models are needed that allow for the reciprocity of trait-range distributions during radiations that unfold over millions of years.

Event-based phylogenetic models have pivotally advanced our understanding of trait and range dynamics of lineages through time (e.g, Butler and King 2004; Ree et al. 2008; Lemey et al. 2010; Goldberg et al. 2011; Uyeda and Harmon 2014; Gill et al. 2017). Standard phylogenetic models, however, generally disregard one or several features that are essential to an idealized model of trait-range evolution. Two key features are (1) that lineages should evolve interdependently with one another and (2) that trait dynamics and range dynamics should be capable of influencing one another. Addressing the first challenge, Nuismer and Harmon (2015) derived a stochastic differential equation (SDE) to test for the effect of biotic interactions under a phylogenetic tree and present day species data (see also Clarke et al. 2017). Because species must be in sympatry to interact, Drury et al. (2016) and Clarke et al. (2017) extended the framework to limit species interactions to those times when lineages were estimated to be in sympatry. Drury et al.’s and Clarke et al.’s methods relies on pre-estimating a distribution of ancestral ranges, and then conditioning on those histories to estimate ancestral trait dynamics. One consequence of this is that the range dynamics unidirectionally influence trait evolution. The second challenge relates to how multiple traits within a single lineage co-evolve. For discrete traits, Sukumaran and Knowles (2018) proposed a joint dependence between geographical and binary traits in a discrete setting by treating the two traits as a single compound trait, then modeling the evolution of that trait with an appropriately structured rate matrix. Lartillot and Poujol (2011) introduced a phylogenetic method that jointly models the co-evolution of continuous traits, discrete traits, and (hidden) lineage-specific evolutionary rates or parameters. And while Lartillot and Poujol’s software implementation of the method, `coevol`, is specialized to study how molecular substitution processes are unidirectionally shaped by life history traits, the underlying design of `coevol`’s inference machinery is suited to more general problems in which continuous traits influence the instantaneous transition rates for models of discrete trait evolution. This is to say that fitting phylogenetic models with either interactions between lineages or with interactions between characters are both challenging problems, each in its own right.

In our work, we build upon these pioneering studies to develop a new parametric model to test for the effect of biotic interactions on the interplay between trait evolution and biogeographic history. First, to better reflect theoretical expectations, we reformulate the SDE describing trait evolution such that the pressure from coexisting species is stronger when lineage traits are most similar, and wanes as traits diverge. Second, instead of supplying a pre-estimated distribution of biogeographic histories, we simultaneously infer biogeographic and trait histories to model interdependence among trait evolution, sympatry, dispersal, and biotic interactions. Third, we allow trait evolution to directly affect the colonization and local extinction rates of lineages throughout their biogeographic history. Specifically, the colonization and local extinction rates for a lineage at a given time depend on the trait values of lineages present across the different biogeographic areas. Notably, our generative model allows direct examination of the distinct contributions of *pre*- and *post*-sympatric niche divergence while attaining secondary contact. For instance, a lineage attempting to colonize a given area might be limited by the similarity among its trait value and those from the species in that area (i.e., competitive exclusion), suggesting a role of *pre*-sympatric niche divergence for successful colonization. Conversely, a lineage could readily colonize any area, independent of the trait distribution found there, but be forced to change because of strong *in situ* interspecific competition, indicating *post*-sympatric niche divergence. We note, however, that we do not model the intricacies of geographic speciation at the nodes and assume that allopatric speciation does not occur; we leave the modeling of this important speciational process to forthcoming work.

Our method fits the model using data augmentation within a Bayesian framework to perform parameter inference, enabling accurate propagation of uncertainty in the posterior distributions by integrating over all trait and biogeographic scenarios found likely by the model. This algorithm has the added advantage of returning joint posterior reconstructions of trait and biogeographic histories, which can be used in post hoc analyses and visualizations. To assess the behavior of our model and to validate our method, we first measure how well it fits a variety of datasets that were simulated under a breadth of evolutionary scenarios. Subsequently, we fit the model to the adaptive radiation of Galápagos finches, an evolutionary system that has been instrumental in exploring phenomena including character displacement, competitive exclusion, and local extirpation due to competition pressure (Lack 1947; Schluter et al. 1985; Grant and Grant 2006). Although our present work focuses on the reciprocal evolution of continuous-valued ecological traits and discrete-valued ranges within and between lineages, our inference framework is extensible to more general models of co-evolution than studied here.

To our knowledge, this is the first study that models biogeographic history and continuous trait evolution as interdependent with one another. This allows assaying previously untestable hypotheses explaining the biogeographic history of clades at the intersection of evolutionary biology and ecology.

## MODEL

### Current approaches for interdependent trait evolution between lineages

Nuismer and Harmon (2015) introduced a continuous trait model where traits of lineages depend on traits of other contemporaneous lineages, allowing biotic interactions among lineages to drive trait divergence and convergence. We follow their derivation of the model, but note that we have modified the notation for the following equations to match analogous parameters in our model. Under the assumption that all lineages are able to interact with each other at any given time (i.e., all are sympatric), weak natural selection and fixed additive genetic variance and population sizes, the change in population mean phenotype for species *i* is given by the following Stochastic Differential Equation (SDE; Eq. S38 in Nuismer and Harmon 2015)

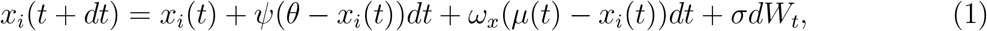

where *ψ* represents the strength of selection, *θ* the selective optimum, *ω*_*x*_ the strength and directionality of competitive interactions, *μ* the expected value of mean phenotypes among all species, *σ* the diffusion rate, and *W*_*t*_ the Wiener process (i.e., standard Brownian motion of Gaussian increments with mean 0 and variance 1). This model couples genetic drift and stabilizing selection (i.e., single-peak Ornstein-Uhlenbeck) with competitive co-evolutionary dynamics; when *ψ* = 0, the model collapses to a random drift with competitive interactions; if, additionally, *ω*_*x*_ = 0, the model becomes a Brownian motion. Lastly, when *ω*_*x*_ ≤ 0, species traits are repelled from a shared average; when *ω*_*x*_ > 0, species traits converge to this average.

The above model assumes that all species in the phylogenetic tree have been sympatric along their evolutionary history, which is often not the case. Drury et al. (2016) expanded on this competition model to incorporate a sympatry matrix among lineages through time. The sympatry matrix effectively limits any interspecific effects upon trait evolution to only those lineages in sympatry at a given time. To do so, let *A*(*t*) represent a time-varying sympatry matrix where entry A_*i,j*_(*t*) = 1 if species *i* and *j* are sympatric at time *t* and 0 otherwise. Then, the change in trait value is given by the following SDE

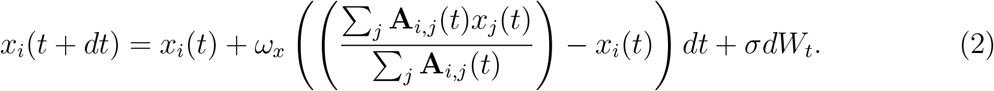

Note, because of non-identifiability, the model assumes no directional selection (*ψ* = 0; Drury et al. 2016). The likelihood of the parameters of interest, *ω*_*x*_, *σ*, and the ancestral state estimate of the MRCA, is a Multivariate Normal density with mean equal to the MRCA state and the scalar product of *σ* with the resulting variance-covariance matrix (Manceau et al. 2017). Drury et al. (2016) derived the SDEs governing the expected variance-covariance through time, and use numerical integration to solve from the root to the tips.

Clarke et al. (2017) proposed a different SDE where species phenotypes are assumed to have normal distributions that phenotypically displace one another in trait space based on their degree of overlap.

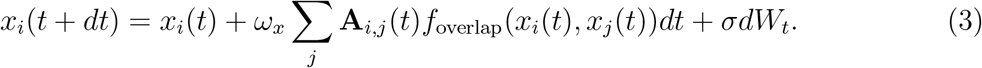

This equation has the advantage of summing over the relative repelling forces from each sympatric species to determine trait evolution instead than just being driven by a community average (Clarke et al. 2017).

One concern with these (and similar) approaches is that biogeographic history is inferred separately from trait evolutionary dynamics, and then conditioned upon when estimating a competition effect on trait evolution. Biologically, the distribution of species traits across areas is likely to directly affect dispersal patterns of lineages along their biogeographic history. For example, extirpation rates might increase among competing lineages while in sympatry, and dispersal rates might decrease for lineages attempting to colonize areas occupied by competitors. More subtly, sequential inference schemes that uniformly average over posterior samples often do not properly weigh the probability of each “upstream” sample when aggregating results under the “downstream” model. This forces the support for each upstream sample to be taken as equal under the downstream model even when that is not true, resulting in the incorrect propagation of uncertainty in species ranges – i.e. a range that is unlikely to be sampled under the trait model would be awarded too much support. Jointly modeling trait and range evolution would circumvent both of these issues, as we describe below.

### Mutually dependent trait and range evolution model

#### Hypotheses framework

There are three parameters that regulate the effect of biotic interactions in our model. The magnitude and directionality of these parameters explicitly examine three expected processes in which interspecific biotic interactions shape biogeographic and trait evolution (Figure 1).

i. Sympatric competition driving character change is described by *ω*_*x*_ (i.e., *post*-sympatry effect of biotic interactions on trait evolution). If *ω*_*x*_ < 0 or *ω*_*x*_ > 0, biotic interactions are driving character divergence and convergence, respectively. If *ω*_*x*_ = 0, no effect of biotic interactions is found when in sympatry, and the particular trait follows a random walk.
ii. The effect of biotic interactions on successful colonization is regulated by *ω*_1_ (i.e., *pre*-sympatric effect of biotic interactions). If *ω*_1_ < 0, lineages have lower rates of successful colonization for areas inhabited by similar species, indicative of competitive exclusion. If *ω*_1_ > 0, lineages have higher rates of successful colonization for areas inhabited by similar species, presumably because of environmental filtering. Evidently, if *ω*_1_ = 0, there is no effect of biotic interactions on rates of colonization.
iii. Finally, *ω*_0_ describes the effect of biotic interactions on rates of local extinction. If *ω*_0_ > 0, more divergent lineages within an area are less likely to go locally extinct, suggesting that competition pressure drives population extirpation. If *ω*_0_ < 0, phenotypically similar lineages within an area are less likely to go extinct, indicative of environmental filtering. Again, if *ω*_0_ = 0, there is no effect of biotic interactions on local extinction rates.

**Figure 1:**
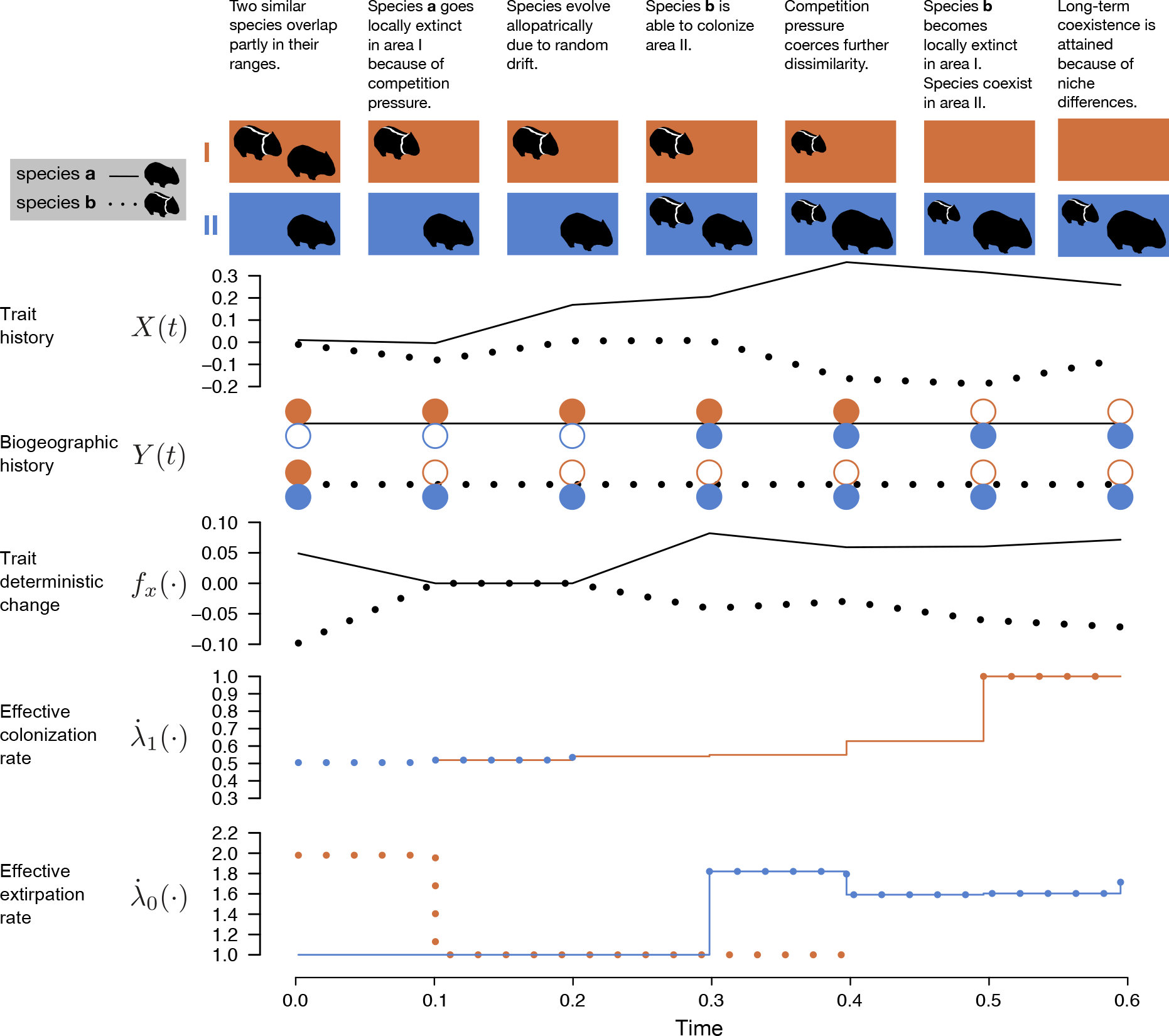
Hypothetical example of a time-discrete history with interdependence between biogeographic and trait evolution for two species, **a**(no stripes and solid lines) and **b**(white stripes and dotted lines), across two areas, **I**(orange) and **II**(blue). We assume that there is *in situ* competition, fixing *ω*_*x*_ = −1, that there is competitive exclusion by fixing *ω*_1_ = −1, and that there is extinction mediated competition by fixing *ω*_0_ = 1. Furthemore, we assume that the random drift *σ*^2^ = 0.1, the base rate of colonization *λ*_1_ = 1 and the base rate of extinction *λ*_0_ = 1. The trait under consideration is the standardized size, specified by *X*(*t*). *Y* (*t*) conveys the specific biogeographic history for each species; filled circles represent the species occupies the area while empty ones that it is absent. The deterministic component of our Stochastic Differential Equation is given by *f*_*x*_(·) and determines the directionality of trait change when in sympatry (Equation 4). Effective rates of colonization per species per area is given by 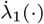; the highest rate of colonization is *λ*_1_ and is given when an area is empty (e.g., last two time steps for area **II**; Equation 5). Effective rates of extinction per species per area is given by 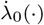; the lowest rate of local extinction is *λ*_0_ and is given when the species is alone in an area (Equation 5). Drawings and values are mathematically consistent following our model.

Table 1 summarizes the effect of model parameters upon the evolution of sympatric lineages for reference.

Adopting a Bayesian perspective allows one to directly detect the effect of sympatric interactions on trait and range evolution. When the 95% highest posterior density (HPD) does not contain the value *ω*_*x*_ = 0, we reject the hypothesis that traits evolve independently among lineages. Similarly, we interpret HPDs that do not contain *ω*_1_ = 0 or *ω*_0_ = 0 as evidence against colonization and extirpation rates being independent of interspecific effects.

#### Model details

We define a joint probabilistic model where rates of area gain and loss for a species may depend on the trait values of all species present in the determined area, and trait values may depend on the trait values of sympatric species (Figure 1). Given a fixed, fully bifurcating and time-calibrated phylogenetic tree with *n* extant species, which we assume as the true tree, and observed data at the tips, we model the biogeographic and trait evolution across time. The crown age of the tree occurs at time 0, with time progressing forward until observing the present values at the tips at time *T*. We denote the entire trait evolutionary history along the phylogenetic tree as *X* and the entire biogeographic history as *Y*. As above, let *x*_*i*_(*t*) be the trait value, in continuous space, for lineage *i* at time *t*. For a set of *K* discrete areas, *k* ∈ {1, …, *K*}, let *y*_*i,k*_ (*t*) be 1 if lineage *i* is present in area *k* or 0 if it is absent at time *t*. Thus, the geographic range of lineage *i* at time *t* can be represented by the vector **y**_*i*_(*t*) = {*y*_*i*,1_(*t*), …, *y*_*i,K*_(*t*)}. Excluding distributions in which species are absent at all areas (i.e., forbidding lineages from going globally extinct), this yields a biogeographic state space containing 2^*K*^ − 1 possible ranges. We sample *n* tips at the present, each with trait value, *x*_*i*_(*T*), and occurring at a subset of discrete locations, **y**_*i*_(*T*). These observations are the result of trait evolution and of species changing their geographic range either by colonizing (area gain) or going locally extinct (area loss) across time.

**Table 1:** Effect of model parameters upon the evolution of sympatric lineages. Trait evolution (*ω*_*x*_) and extirpation (*ω*_0_) parameters are informed by sympatric differences in traits in the currently inhabited area(s). The colonization parameter (*ω*_1_) is informed by differences in traits between the colonizing lineage and the resident trait distribution in the area to be colonized.

| Parameter | Value | Effect of sympatry |
| --- | --- | --- |
| $\omega_x$ | 0 | No effect |
| | $< 0$ | Traits diverge |
| | $> 0$ | Traits converge |
| $\omega_1$ | 0 | No effect |
| | $< 0$ | Lower colonization rates |
| | $> 0$ | Higher colonization rates |
| $\omega_0$ | 0 | No effect |
| | $< 0$ | Lower extirpation rates |
| | $> 0$ | Higher extirpation rates |

We model the effect of competition on the trait evolution of lineage *i* using the following SDE

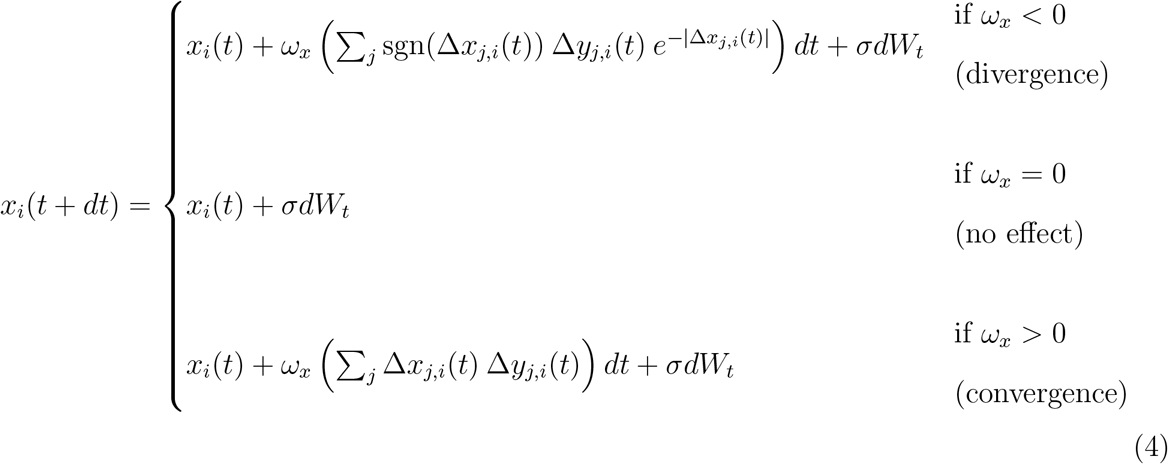

where sgn(*x*) = {−1 if *x* < 0, 0 if *x* = 0, and 1 if *x* > 0}, and

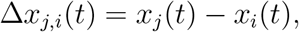

and

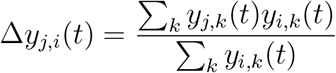

represent trait and range differences between lineages, respectively. That is, the strength of biotic interactions for the focal lineage *i* at time *t* is measured in relation to the weighted sum of trait differences with other species, Δ*x*_*j,i*_(*t*), scaled proportionally to the amount of range overlap, Δ*y*_*j,i*_(*t*). Figure 2a illustrates the behavior of this SDE. Importantly, it befits the theoretical expectation that competition strength should wither as trait dissimilarity increases. Fortunately, the inference scheme that we use (see below) provides great flexibility in specifying the deterministic part of the SDE, as long as it is a function of the form *x*_*i*_(*t* + *dt*) = *f*_*x*_(*X*(*t*), *Y* (*t*), *ω*_*x*_, *dt*) + *σdW*_*t*_.

**Figure 2:**
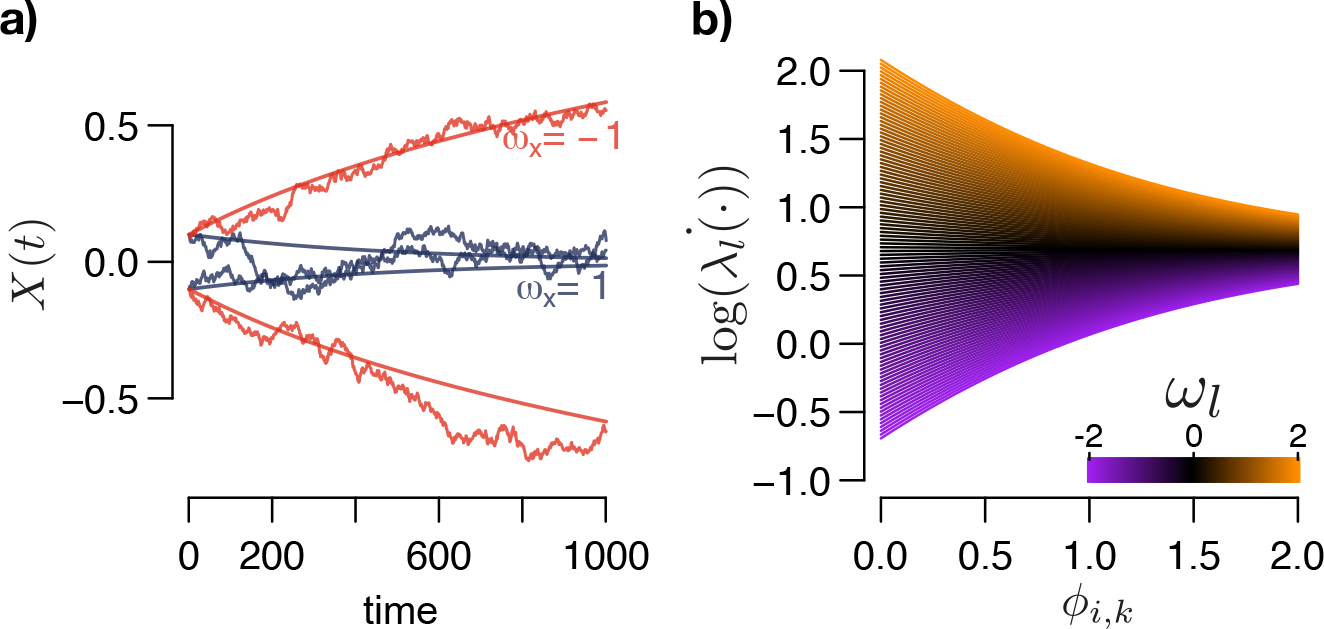
Functional forms for the joint evolution of trait and ranges. **a)** An illustration of the Stochastic Differential Equation (SDE) used to model the role of biotic interactions in trait evolution. We plot trait evolution as the stochastic (diffusion) component superimposed upon the deterministic (interspecific) component. At time *t* = 0, the phenotypic values of two lineages, *X*_*a*_(*t*) = −0.1 and *X*_*b*_(*t*) = 0.1, evolve according to the *in situ* biotic interations parameter, *ω*_*x*_. If *ω*_*x*_ < 0, the lineages repel each other, if *ω*_*x*_ = 0, the lineage evolves by random drift, and if *ω*_*x*_ > 0, they attract each other. **b)** Functional form relating trait differences for lineage *i* and those in area *k*, *ϕ*_*i,k*_, and the logarithm of the effective rates of colonization or extinction, 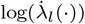. Here, *l* indicates a gain (1) or loss (0) event, for different values of *ω_l_*. Purple colors represent *ω*_*l*_ values close to −2 and orange colors close to 2. If *ω*_*l*_ < 0, lower trait differences between lineages suffer higher penalties in rates of colonization or extirpation relative to larger differences, if *ω*_*l*_ = 0, then 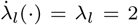, and finally if *ω*_*l*_ < 0, larger trait differences between lineages enhance the rates of colonization or extirpation.

To test the effect of biotic interactions on biogeographic history, we allow for rates of colonization and local extinction for a given lineage *i* to vary according to the similarity between its phenotype *x*_*i*_ and that amongst all species currently in an area. Specifically, let **u**, **v** be geographic ranges that differ only on area *k*, with *u*_*k*_ = 0 and *v*_*k*_ = 1, and let 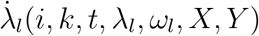 for *l* = {0, 1} be the instantaneous rates of area gain or loss, respectively, for area *k* and lineage *i* at time *t*. Then, we define

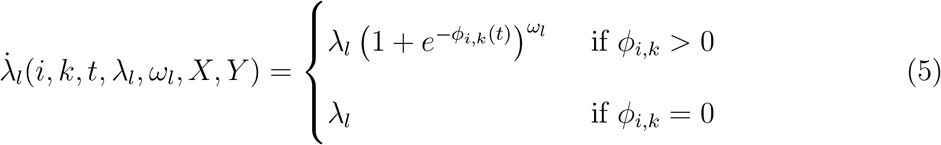

Where

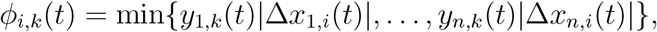

*λ*_*l*_ is the “basal” rate of colonization or extinction, *ω*_*l*_ describes the effect of biotic interactions on rates of colonization or extirpation, and *ϕ*_*i,k*_ (*t*) is the minimal distance in trait space between lineage *i* and those in area *k*.

Equation 5 is a simplified version of the Generalized Logistic function (see Appendix). Note that when *ω*_1_ is negative, these functional forms designate *λ*_1_ as the maximum colonization rate when an area is unoccupied, and the presence of other species induces a penalty on the rates, in turn, when *ω*_1_ is positive colonization rates are enhanced. Similarly, *λ*_0_ is the rate when an area is unoccupied, and sympatric species induce a rate increase with *ω*_0_ > 0 and a decrease when *ω*_0_ < 0. In both cases, the penalty is dependent on the minimum distance between the focal species *i* and those in the area being considered *k* (i.e., *ϕ*_*i,k*_). Thus, the magnitude of *ω*_1_ and *ω*_0_ reflect the relative effect in which biotic interactions affect biogeographic rates (Figure 2b).

#### A discretized time scheme

We wish to compute the probability of a single, exact co-evolutionary history of traits and ranges along all branches of a phylogeny. Even for a single trait-range history, we were unable to derive an analytical form of the transition probabilities for trait evolution (Eq. 4) and range evolution (Eq. 5) as functions of continuous time. Thus, following Horvilleur and Lartillot (2014), we represent the continuous-time processes of trait and range evolution in discrete time. This time discretization serves two purposes: first, it lets us derive the discrete-time transition probabilities we need to compute the model probability; and, second, it provides a basis to rapidly query the complete evolutionary state shared across lineages, areas, and traits at regular time intervals, which is essential for computing the transition probabilities.

Figures 3a and 3b illustrate an example output of our two-stage discretization procedure, which results in the ordered vector of times, ***τ***. The procedure works as follows. Let *t*_0_ = 0 be the crown age of the tree, and let *T* be the time at which we observe the tip trait values, *X*_obs_, and range values, *Y*_obs_. Also, let branch *b* have a start time *t*_*bs*_ and end time *t*_*bf*_, such that *t*_*b*_ = *t*_*bf*_ − *t*_*bs*_. The first stage divides each *t*_*b*_ into *K* + 1 equally spaced time slices (i.e., the number of areas plus one), yielding the vector of sampling times ***τ***_*b*_ = {*t*_*bs*_ = *τ*_*b*, 1_, …, *τ*_*b,K*+2_*, t*_*bf*_ = *τ*_*b,K*+3_}. Because we only allow one event per time step, the number of slices, *K* + 3, guarantees that lineage *i* has more than the minimum number of steps possibly needed to evolve from range **y**_*i*_(*b*_*s*_) to **y**_*i*_(*b*_*f*_) in the case where **y**_*i*_(*b*_*s*_) is absolutely different from **y**_*i*_(*b*_*f*_) (for example, it would take at least three events for the range **y**_*i*_(*b*_*s*_) = {0, 0, 1} to evolve into range **y**_*i*_(*b*_*f*_) = {1, 1, 0}). The second stage sets a minimum time step allowed in the analyses, *δt*_min_, and proceeds forwards in time to subdivide the remaining periods such that no time step is larger than *δt*_min_. In practice, we standardize *δt*_min_ using the percentage of the tree height for comparability. This procedure results in a sorted vector of sampling times ***τ*** = {*t*_0_ = 0, …, *T*} that are shared among all contemporaneous lineages throughout the clade’s history. For each branch *b* with sampling times ***τ***_*b*_ ⊆ ***τ***, we end up with a time ordered set describing the trait evolution of the lineage, 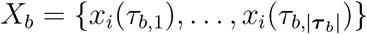. Likewise, for each branch *b*, we record an ordered set of vectors describing the biogeographic history of the lineage, 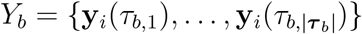.

**Figure 3:**
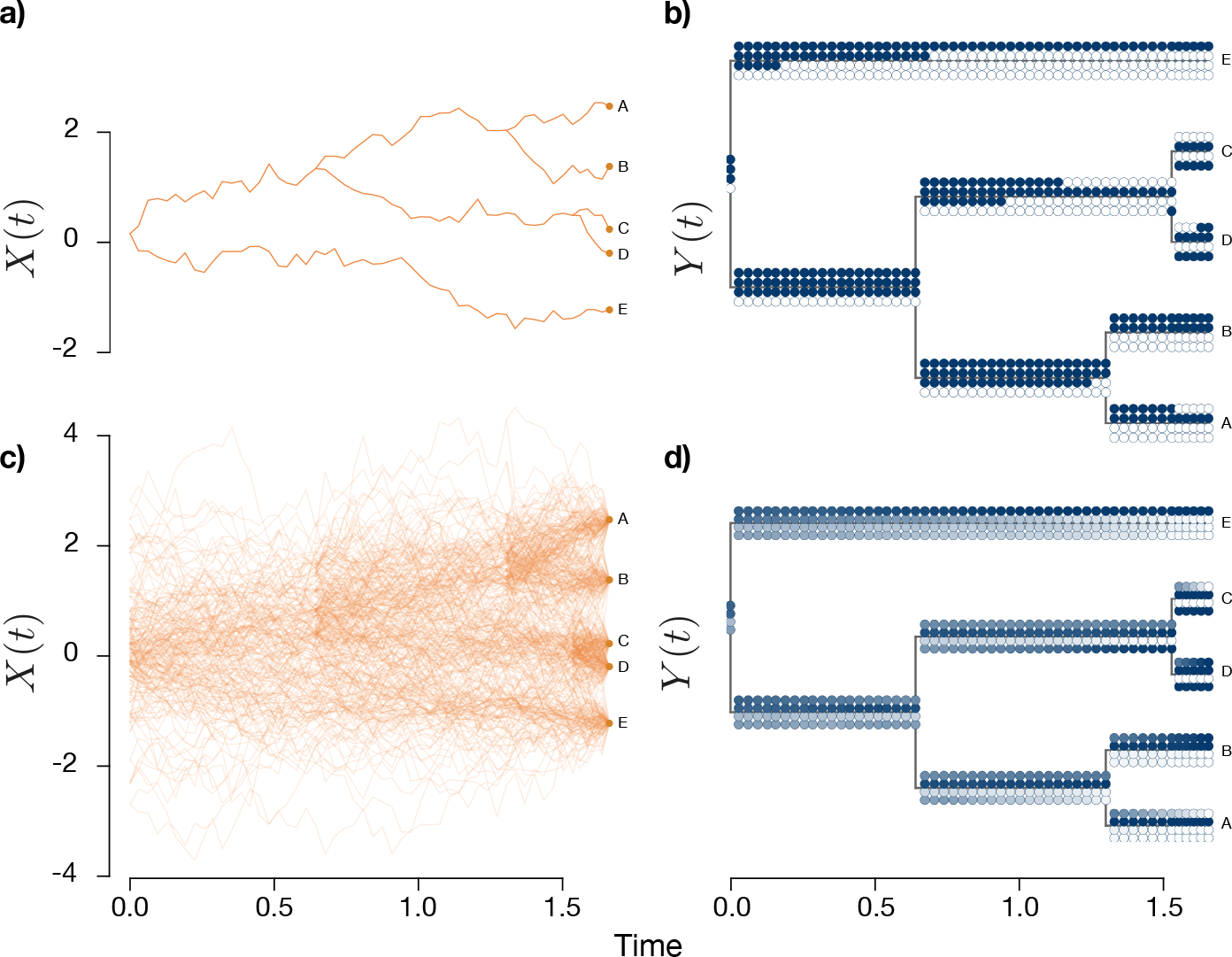
Illustration of the discretized data augmentation used from a simulation performed on an ultrametric tree of 5 tips and four areas with in situ competition (i.e., *ω*_*x*_ = −1). **a)** One random sample trait history, *X*(*t*), from the posterior. **b)** One random sample of biogeographic range history, *Y* (*t*), from the posterior across four areas. Each time sample has four circles in vertical orientation, each representing one of the areas. Filled circles represent occupied areas while empty circles represent absence. Note that all branches have at least five internal discrete sampling times, that is, one more than the number of areas in the current system. We set the minimum time interval here to be 2% for the tree height for illustration purposes. **c)** Marginal posterior data augmented histories based on 100 samples in trait with translucency. **d)** Corresponding marginal biogeographic histories. Darker tones represent higher marginal probabilities of area occupancy.

#### Likelihood calculation

We are not aware of an analytical form for the transition probabilities corresponding to the range-dependent trait evolution model (Eq. 4), so we approximate the likelihood using the Euler-Maruyama method (see Appendix). The likelihood for trait evolution for branch *b* is then

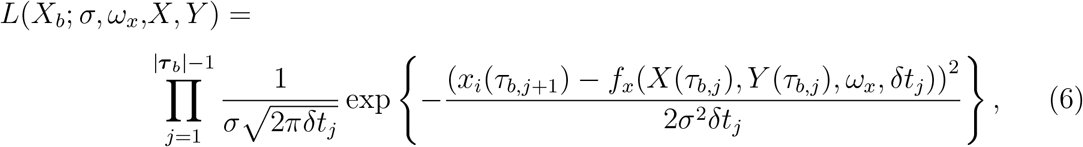

where *δt*_*j*_ = *τ*_*b,j*+1_ − *τ*_*b,j*_.

The likelihood for the biogeographic history in discrete time can be deconstructed into a series of events and nonevents within small windows of time. An event is defined as either an area colonization or loss, and a nonevent as no change in state. Let *l* = {0, 1}, then the likelihood after some time *δt* for area *k* is

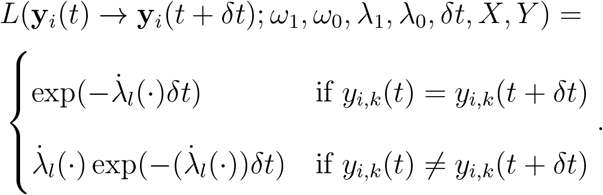

Then, the likelihood for branch *b* across all areas is:

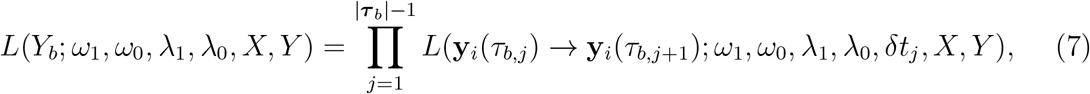

where *δt*_*j*_ = *τ*_*b,j*+1_ − *τ*_*b,j*_.

The prior probabilities for each state are usually set to the stationary frequencies given by the dispersal rates *λ*_1_ and *λ*_0_. We could not derive an analytical solution for these frequencies, so we add a long branch (twice the tree height by default) at the root and simulate geographic range evolution to approximate geographic range frequencies at the root (Landis et al. 2013). Under the model assumptions, there is no trait data for the stem branch (and, given the tree, there is no competition since only one lineage of the clade is alive), so the likelihood computation can be done in continuous time. Let 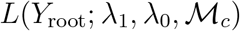 denote this likelihood and ***θ*** = {*σ*, *ω*_*x*_, *ω*_1_, *ω*_0_, *λ*_1_, *λ*_0_}. Then, by incorporating the trait evolution likelihood and multiplying across all branches, we get the following joint likelihood:

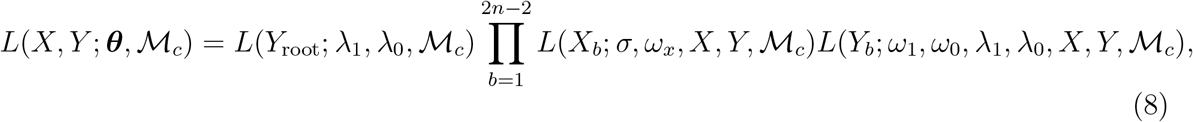

where 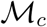 is the model incorporating biotic interactions.

#### Collision probability

It is possible that a species range gains and then loses an area (or vice versa) so rapidly under the idealized continuous-time model that those events would go undetected by our discrete-time model. Such “collisions” of events within a single discrete time bin might lead to underestimating the area colonization and loss rates. We estimate an upper bound on the collision probability, 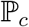 that two or more range evolution events occur within a fixed *δt*, such that our sampling would not detect them. Specifically, let *δt* be a time interval for which we sample Y and X at the beginning, *t*_*s*_, and at the end, *t*_*f*_, where *t*_*f*_ = *t*_*s*_ + *δt*. If the lineage is present in area *k* at time *t*_*s*_, the lineage could lose this area and regain it before we are able to register such event in *t*_*f*_. Let *r* = (*λ*_1_ + *λ*_0_)*δt*, then the probability that two or more events at times occur within *δt* is

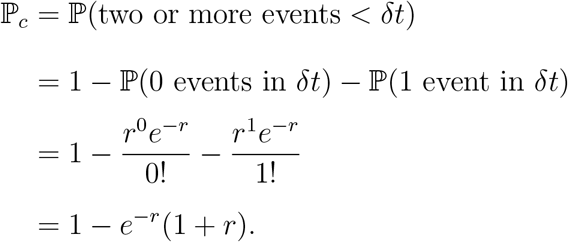

We consider *δt* to be the largest interval in the analysis, thus providing a somewhat conservative measure of collision probability (given that there are smaller intervals following our discretization procedure). However, since the actual rates rely on the specific interaction between trait value differences and *ω*_1_ and *ω*_0_, this measure does not necessarily reflect the actual collision probability, yet it still is a source of objective information on amount of approximation error. We monitor 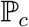 during inference to provide a measure of error given the particular parameters and defined *δt*.

### Markov Chain Monte Carlo with data augmentation

The main impediment when inferring under such a joint model is the mutual dependence of the trait evolutionary history, *X*, and the biogeographic history, *Y*. At any given time, trait evolution for one species depends on the traits of those species it is sympatric with, and the set of species that are able to coexist in sympatry is contingent on the concurrent trait distribution. This, in part, renders common inference procedures such as the derivation of an analytic solution or pseudo-exact likelihood by numerical integration of SDEs infeasible. Rather than analytically integrating over all possible evolutionary histories, we use data augmentation (DA) to numerically sample over those histories (Robinson et al. 2003; Landis et al. 2013). Under DA, one repeatedly simulates otherwise unobservable data to evaluate the probability of the parameters ***θ*** under both the observed data *D*_obs_ and the augmented data *D*_aug_. Among several advantages of using DA is the fact that, for certain problems, simpler and more efficient likelihood functions exist when augmented data is generated. By repeatedly proposing different realizations of *D*_aug_ across the MCMC, one numerically averages over the augmented data to obtain the joint posterior of evolutionary histories and model parameters, 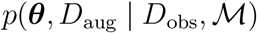. In particular, we are interested in computing the posterior probability of all the parameters given the observed data. The posterior probability of one single biogeographic and trait history is

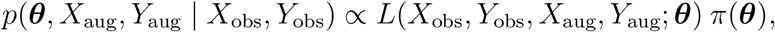

where *π* is the prior distribution of ***θ***. We describe the initialization procedure for *X*_aug_ and *Y*_aug_ in the Appendix. Figure 3c,d shows a sample from the marginal posterior for DA trait and biogeographic histories from a simple simulation. We sample augmented evolutionary histories and evolutionary parameters using the Metropolis-Hastings algorithm (Metropolis et al. 1953; Hastings 1970).

### Parameter, trait history, and range history proposals

Standard slide and scale moves are used to proposed new parameter values for *σ*, *λ*_0_, *λ*_1_, *ω*_0_, *ω*_1_, and *ω*_*x*_ (see Appendix).

We generate proposals for the trait history, *X*_aug_, by adding a Gaussian deviation to a uniformly sampled *x*_*i*_(*t*), such that *x*_*i*_(*t*)′ = *x*_*i*_(*t*) + N(0, *s*), where *s* represents the tuning parameter. The acceptance ratio for this proposal is

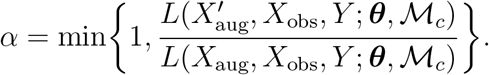

In addition, we generate less conservative updates by proposing branch-wide updates for *X*_aug_. We use random samples from an independent distribution for *σ*\* to generate Brownian bridges for branches in the tree (details for generating a Brownian bridge are given in the Appendix). First, we sample *σ*\* ~ Lognormal(0, 1) and a branch uniformly and generate a Brownian bridge holding the end nodes constant. Similarly, following Horvilleur and Lartillot (2014), we sample an internal node uniformly and generate a new node state under Brownian motion and generate Brownian bridges for the three adjoining branches. The acceptance ratio for the these proposals is

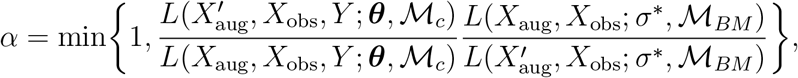

where 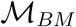 denotes the Brownian Motion model.

To update the range history, *Y*_aug_, we select an internal node uniformly at random, including the root, and sample a new geographic range from the joint density under the mutual-independence model, 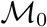. We use random samples from an independent distribution for 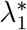 and 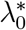 to generate DA biogeographic histories under 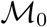. We improve efficiency and acceptance rates of biogeographic histories by disallowing colonization and extirpation rates to be too dissimilar. Therefore, we randomly sample *υ* ~ Lognormal(0, 1), and then multiply *υ* by a Lognormal distribution with expectation of 1 and low variance such that 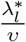 ~ Lognormal(−0.044, 0.3) for *l* ∈ {0, 1}. Using the rejection sampling described in Landis et al. (2013) and the Appendix, we then sample new biogeographic histories along the three adjoining branches such that they are consistent with the new sampled geographic range at the node and those at the end nodes. The simplified Metropolis-Hastings acceptance ratio (*α*) for this proposal is

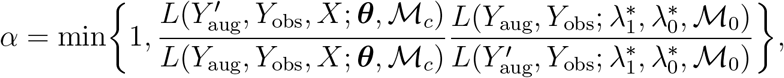

where the first term is the ratio between the likelihoods of the proposed and current biogeographic histories under the full model, 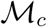, and the second term is the proposal density ratio under the mutual-independence model, 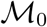. Additionally, we perform more moderate proposals for range evolution by mapping biogeographic histories on a branch sampled at random, leaving the end nodes constant. The acceptance ratio for this branch update is the same as for the node update. As mentioned earlier, daughter lineages inherit the same geographic range as their parent lineage at speciation times. This mimics a very particular case of sympatric speciation, a strong assumption for the biogeographic history of some clades. The intricacies of geographical speciation will be left for future work (e.g., Ree et al. 2005; Matzke 2014).

Finally, to better explore parameter space, we make joint *X*_aug_ and *Y*_aug_ proposal updates. For the first joint update, we uniformly sample a branch and update the trait history using a Brownian bridge proposal and update biogeographic history using stochastic mapping as described above. Secondly, we uniformly sample an internal node and generate a joint proposal for the node and the the three adjoining branches. The acceptance ratio for these proposals is

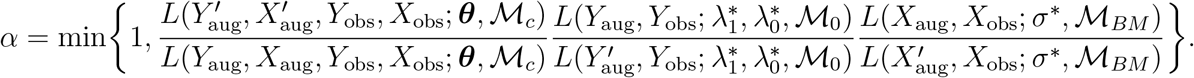

#### Software

We denote this model as “TRIBE” (which stands for “Trait and Range Interspecific Biogeographic Evolution”) and implement it in a new open source package named “Tapestree” (https://github.com/ignacioq/Tapestree.jl) that we wrote in Julia (Bezanson et al. 2017). This software makes available the `tribe()` function for inference and the `simulate tribe()` for simulations given a fixed tree. We note that, in the software, we allow the user to fix to 0 any or all of the parameters governing the effect of biotic interactions (i.e., *ω*_*x*_, *ω*_0_, & *ω*_1_).

### Simulations

We use simulations to explore model behavior. To simulate biogeographic histories under this model, we take advantage of the following approximation. Let *V* be a random variable denoting the time of an event and *λ*(*t*) be the event rate at time *t*, then given a small enough time step *δt*, we have

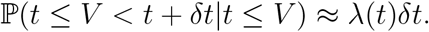

Thus for a given lineage and timepoint, we use the above time step size for all areas across the geographic range. If there is more than one event within one time step as defined by our time discretization scheme, we reject and sample again. Similarly, to simulate trait evolution under the competition model, we, again, take advantage of the Euler-Murayama method detailed in the Appendix. Simulation code, given a phylogenetic tree, can be found at https://github.com/ignacioq/Tapestree.jl.

We simulated phylogenetic trees using a pure-birth process until reaching 25 species and set the MRCA trait value to 0 and the number of areas to 12. Given the relatively large parameter space, we used the same values for *λ*_1_, *λ*_0_ and *σ*^2^ across all simulations, and explored different combinations of the parameters regulating the biotic interactions. In particular we simulated 10 different scenarios with *λ*_1_ = 1.0, *λ*_0_ = 0.4 and *σ*^2^ = 0.16, and the following combinations of (*ω*_*x*_, *ω*_*1*_, *ω*_0_): (0, 0, 0), (−2, −2, 0), (−2, 2, 2), (−2, 0, 0), (2, 0, 0), (2, −2, 2), (2, 0, 0), (0, −2, 2), (0, −2, 0), and (0, 0, 2). Each scenario was simulated 100 times to yield a total of 1000 simulations. While not exhaustive, these simulations allow us to test the power and bias of our model with regard to each of these three parameters. Further exploration of parameter space is encouraged for the future.

We ran MCMC inference on each simulation for 100, 000 iterations, logging every 100^th^ iteration, discarding the first 50, 000 samples obtained during the adaptive burn-in phase. We note that each iteration corresponds to > 55, 000 parameter updates (the user can adjust the weights for each parameter). We used ambiguous priors for all parameters, specifically, we used a normal prior of mean 0 and standard deviation of 10 for *ω*_*x*_, *ω*_1_, and *ω*_0_, and an exponential prior of mean 10 for *σ*^2^, *λ*_1_ and *λ*_0_. Most of the effective sample sizes (ESS) for all parameter in each simulation were > 300, but in a few cases *σ*^2^ or *ω*_*x*_ had lower ESS; we made sure that the ESS for each parameter was at least > 150.

We perform statistical evaluation using highest probability density intervals (HPD) for all the parameters. Overall, our model is able to recover most of the simulated parameter values and associated uncertainty. The posterior median estimates reflect the simulated values (Figure 4) and 95% coverage probability based on HPD for parameters reflecting biotic interactions are over 0.90 for most scenarios (Figure 5). Most importantly, our model is able to reliably discern when there is no effect of biotic interactions for *ω*_*x*_ and *ω*_0_ (Figures 4 & 5). Estimates of the posterior mean of *ω*_*x*_ behave without bias when the true value is negative, yet the have a marginally positive bias towards more positive values when it is ≥ 0; this is most likely because of an increase in skew in the posterior distribution as *ω*_*x*_ increases. The 95% HPD coverage is close to 0.95 for all scenarios (Figure 5).

**Figure 4:**
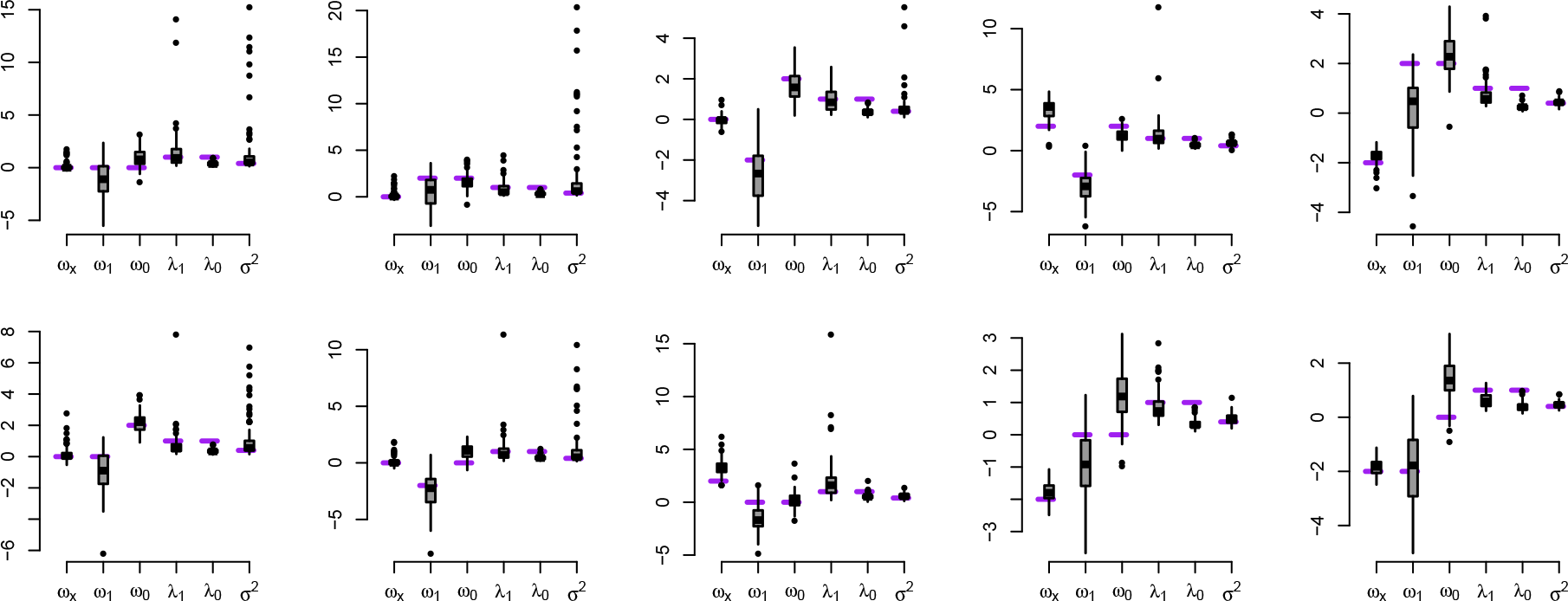
Boxplots of median posterior estimates from the different simulation scenarios. Each panel represents 100 different simulations in pure-birth trees with 25 tips and 10 areas. The true values used for the simulations are represented in horizontal dotted purple lines.

**Figure 5:**
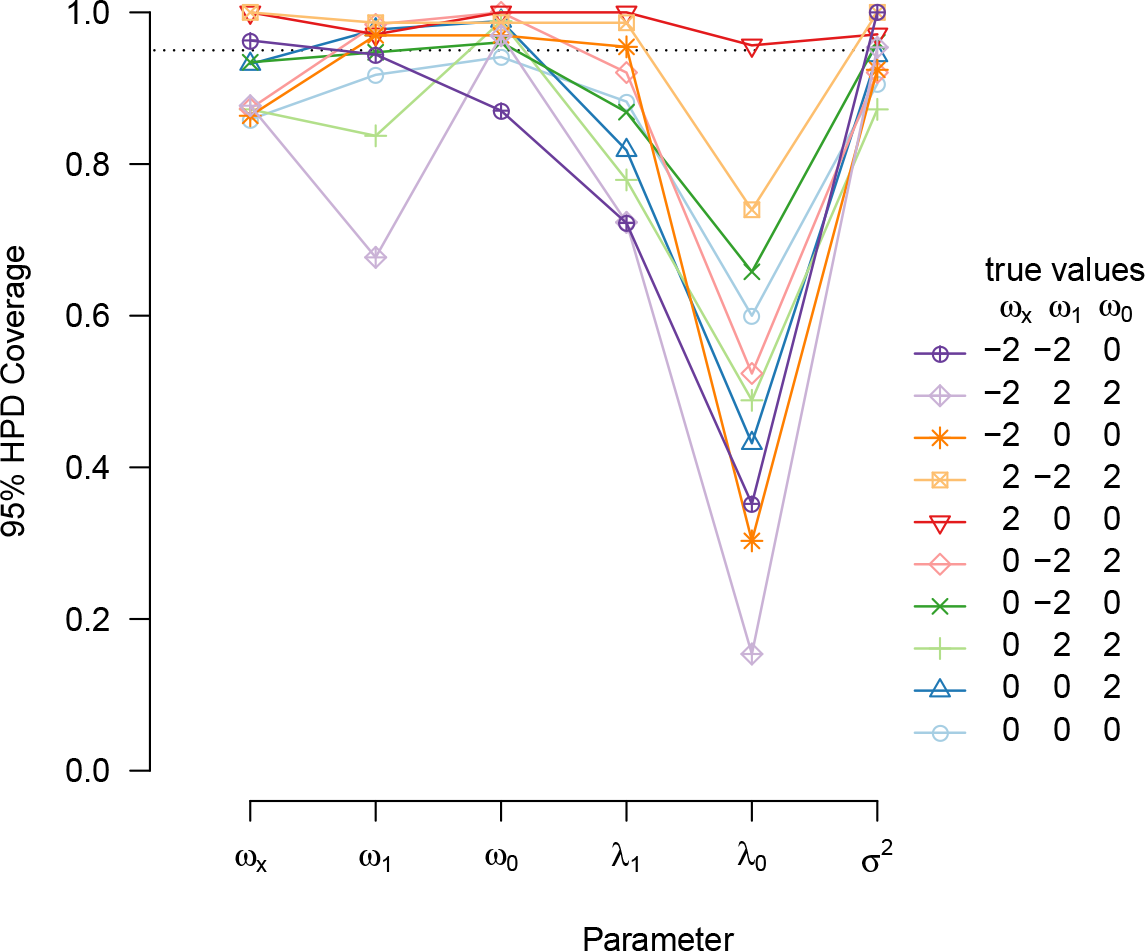
Posterior statistical 95% Highest Posterior Density (HPD) coverage for the 10 simulation scenarios for each parameter. Each symbol and color represents a different set of true values used for the simulation, corresponding to those used in Figure 4. The dotted line corresponds to 95% of HPDs across simulations covering the true simulated parameter.

Nonetheless, we find a minor bias in *ω*_1_, the parameter regulating competition on colonization rates. Recovered values for *ω*_1_ are biased toward lower values, however, the coverage remains at least 90% for scenarios for scenarios with *ω*_1_ = 0, yielding acceptable false positive rates for competition (Figures 4 & 5). We find the greatest bias and lowest coverage for scenarios in which *ω*_1_ > 0, and may result in false negatives for facilitation in colonization rates. Finally, we find that posterior estimates of *λ*_1_, *λ*_0_ are somewhat underestimated, and their medians are usually lower that the simulated value. While concerning, this is likely due to the interaction with the phenotypic traits and does not preclude our ability to make inference on the effect of biotic interactions on biogeographic and phenotypic evolution. Overall, most likely increasing the number of taxa and areas will result in higher power.

#### Impact of δt_min_

To evaluate the impact of different *δt*_min_ in parameter estimates, we performed inference on the same data with five different *δt*_min_ = {0.99, 0.2, 0.1, 0.01, 0.005}. Note that it is often the case that increasing values of *δt*_min_ to be greater than *ca.* 0.2 gives the same discretization scheme and thereby similar results because our discretization procedure minimally includes times for the start, end, and *K* + 1 intermediate time points along every branch in the tree (clearly, this threshold is relative to the structure of the tree). The simulations were conducted with the same pure-birth tree of 25 species and 4 areas, and the following parameter values: *ω*_*x*_ = −2, *ω*_1_ = 1, *ω*_0_ = −1, *σ* = 0.8, *λ*_1_ = 4 and *λ*_0_ = 2. We ran the analysis with an adaptive burn-in of 50000 iterations and a sampling chain of 100000.

We find that the impact of *δt*_min_ has minor consequences on the parameter estimates in the posterior distributions (Supplementary Figure 1). This is most likely due to the discretization procedure that ensures that each branch will be subdivided into at a number of units greater (by one or more) than the number of areas. Such discretization is thus finer towards the tips, where more branches overlap in time, and where inference is less uncertain (since is more proximate in time to the observed trait and biogeographic data). We find *σ*^2^, *ω*_1_ and *ω*_0_ to be marginally affected by the the choice of *δt*_min_. The differences are slightly pronounced in *ω*_*x*_, *λ*_1_, and *λ*_0_, particularly in terms of precision. This is expected as we reiterate that we are approximating the likelihoods, and a finer discretization will be less biased. For instance, a finer discretization allows higher rates of colonization and extinction to be sampled in the posterior (Supplementary Figure 1). Larger *δt* values between sampling times incur in high collision probabilities, thus ignoring high rates of state changes and setting an upper limit on the inference of rates of state change. Given our simulation results and required computational efficiency, we suggest that a *δt*_min_ = 0.01 yields an acceptable representation of the model likelihood.

### Empirical application: Darwin’s finches in the Galápagos

We use our model to study how biotic interactions have shaped the biogeographic and trait evolution of Darwin’s finches on the Galápagos islands (Grant 1999). We used the species phylogenetic tree from (Lamichhaney et al. 2015) for 14 species and obtained corresponding breeding distributions across the major Galápagos islands (19 islands, including Cocos island), following Table 1.2 in Grant and Grant (2011) and phenotypic measurements from Clarke et al. (2017), originally compiled in Harmon et al. (2010) from which we obtained the data for *Certhidea olivacea*. Specifically, we used three beak measurements: length (culmen), width and depth (gonys), and tarsus and wing length, all with natural logarithmic transformations. Given the high correlation between the three beak measurements, we used the first and second principal components (which together explained > 99.6% of the variance). The first component mostly corresponds to size, while the second corresponds to overall shape (Supplementary Figure 2; Grant and Grant 2002). The finch data used in this study can be found in the Supplementary Table 1. We ran separate models for these four trait values, for 500 thousand iterations with an adaptive burn-in phase of 50 thousand.

**Table 2:** Posterior estimates for Darwin’s finches analysis.

| Trait | Posterior median and HPD estimates |  |  |  |  |  |
| --- | --- | --- | --- | --- | --- | --- |
| | $\omega_x$ | | $\omega_1$ | | $\omega_0$ | |
| Beak size (PC1) | -0.46 | [-1.30, 0.54] | -6.38 | [-9.89, -2.80] | 1.53 | [-0.74, 3.90] |
| Beak shape (PC2) | 1.28 | [0.12, 3.57] | -4.28 | [-8.63, -0.46] | 2.2 | [-0.16, 4.90] |
| Tarsus length | 2.41 | [-0.04, 6.46] | -4.60 | [-6.20, -1.50] | 2.87 | [-0.26, 5.08] |
| Wing length | 0.61 | [-0.10, 5.30] | -4.86 | [-6.86, -1.23] | 0.43 | [-0.99, 3.50] |

We find that *in situ* trait evolution behaves very differently across the four traits studied here (Figure 6). Overall, we do detect a signal of competitive exclusion (*ω*_1_ < 0), with varied levels of strength. Beak morphometrics (the first and second PCA components relating to size and shape, respectively) display different results (Figure 6e). Beak size shows divergence in sympatry (median *ω*_*x*_ = −0.46, 95% HPD = [−1.3, 0.54]); on the other hand beak shape shows convergence (median *ω*_*x*_ = 1.28, 95% HPD = [0.12, 3.57]). These traits display values of *ω*_1_ < 0, suggesting strong signals of competitive exclusion, particularly for beak size (median for size = −6.38 [−9.89, −2.8]; median for shape = −4.28 [−8.63, −0.46]). Finally we find a weak effect of biotic interactions on the influence of beak size and shape on local extirpation (median *ω*_0_ for size = 1.53, [−0.74, 3.9], for shape = 2.2, [−0.16, 4.9]).

**Figure 6:**
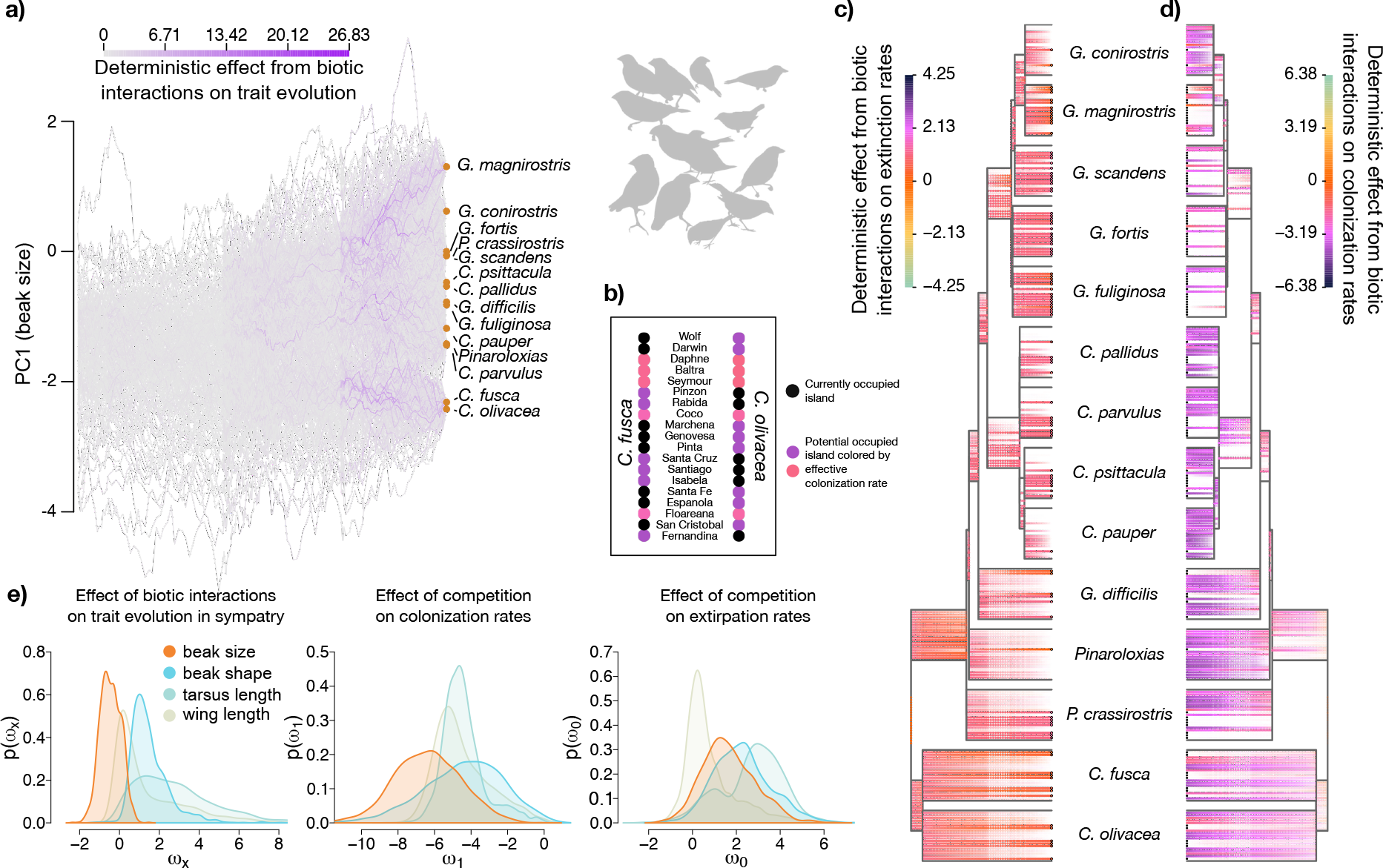
Empirical results for the effect of biotic interactions on the trait and biogeographic evolution of Darwin’s finches. **a)**100 data augmented trait histories for PC1 (beak size). Absolute deterministic effects of biotic interactions on trait evolution for sympatric lineages are colored from grey (isolated evolution under Brownian motion) to purple (strongest effect of biotic interactions). **b)** Example of present-day effect of biotic interactions in colonization rates between two species that are phenotypically similar, *Certhidea fusca* and *C. olivacea*. The areas are displayed as circles arranged in a column, with currently occupied areas (islands) in black and unoccupied areas colored according to effective colonization rates following the color scale in Figure 6d (below). Note that areas occupied by the sister species suffer a colonization penalty and reflect competitive exclusion in beak size as given by our model. **c)** Marginal data augmented biogeographic histories for the same 19 areas shown in Figure 6b. Alpha opacity denotes the marginal probability of presence at a given time for a given lineage-area. The color scale represents the average effect of biotic interactions on local extinction rates (purple denoting higher rates of local extinction and orange, no influence). Currently occupied areas are shown with black unfilled circles at the tips. **d)** As in Figure 6c, but alpha opacity denote the marginal probabilities of absences at a given time for a given lineage-area, and the color scale represent the average effect of biotic interactions on colonization rates (purple denoting lower rates of colonization and orange no influence). Currently occupied areas are shown with black filled circles at the tips. **e)** Posterior marginal densities for the parameter governing biotic interactions (*left: ω_x_*, *middle: ω*_1_, *right: ω*_0_) for each of the four phenotypic traits analyzed separately. The results suggest *in situ* competition for beak size and strong convergence for tarsus and beak shape. All traits show strong penalization for colonization when similar. See text for further details. Finch silhouettes from Caroline O’Donnell, redrawn from Biological Sciences Curriculum Study, *Biological Science: Molecules to Man*, Houghton Mifflin (1963).

Figure 6b focuses on just two finch species that share similar beak sizes at one moment time (present-day), but do not overlap on their geographic distributions. Evidently, *Certhidea fusca* is expected to suffer from lower colonization rates into areas that are occupied by *C. olivacea*, with reciprocal effect for the latter species attempting to colonize areas occupied by the former. This example highlights how our approach may identify whether the allopatric (or sympatric) distribution between species is a product of biotic interactions or independent of them.

We find a signal of *in situ* convergence for tarsus length (Figure 6e), (median *ω*_*x*_ = 2.41, 95% HPD = [−0.04, 6.46]). We observe a strong effect of competition in colonization rates (median *ω*_1_ = −4.6, 95% HPD = [−6.2, −1.5]) and no effect of biotic interactions on extirpation rates (median *ω*_0_ = 2.87, 95% HPD = [−0.26, 5.08]). Biotic interactions had no effect for wing length when in sympatry (median *ω*_*x*_ = 0.61, 95% HPD = [−0.1, 5.3]), but instead find strong competitive exclusion (median *ω*_1_ = −4.86, 95% HPD = [−6.86, −1.23]). We find no evidence for an effect of competition in driving local extinction (median *ω*_0_ = 0.43, 95% HPD = [−0.99, 3.5]). Together, these results suggest that there is strong evidence for competitive exclusion in Darwin’s finches in beak morphology and, particularly, in wing length (Figure 6e). Overall, beak size seems to reflect a key competition axis that has driven trait divergence and shaped biogeographic history.

## Discussion

Ever since Darwin (1859), biologists have strived to understand the extent and generality of different biological processes in driving current patterns of diversity (Simpson 1953; Mayr 1970; Schluter 2000). Building on previous developments, we introduce a simple but extensible model that integrates discrete biogeographic processes with continuous phenotypic evolution, enabling direct tests on those processes underlying trait evolution, biogeographic history, biotic interactions and community assembly.

### Darwin’s finches

We show how biotic interactions influence trait and biogeographic evolution using the radiation of Darwin’s Finches and find that competition has played a role in beak size divergence when different species come into sympatry (Figure 6). While this is in accordance with previous findings (e.g., Lack 1947; Grant and Grant 2006; Clarke et al. 2017), we only find evidence for trait divergence in beak size but not in shape. Instead, our results suggest that bill shape and tarsus length have converged among coexisting species. Presumably, the harsh and unpredictable environmental conditions in the archipelago give rise to strong selection against variants (Price et al. 1984), leading to long term morphological convergence in some traits across the different islands. Indeed, character displacement presupposes that there exists niche space to be displaced into, but extreme events such as droughts severely reduce the number of available sources within an area (Grant and Grant 2011), removing accumulated trait variance. Thus, our results suggest that there is character displacement in beak size but other traits might be phenotypically constrained given the available environment. Future model enhancements could incorporate environmental information to distinguish biotic from abiotic effects. Similarly, persistent introgression during the clade’s evolutionary history could lead to some the observed convergence in morphology (Grant et al. 2004; Grant and Grant 2008; Farrington et al. 2014; Lamichhaney et al. 2018).

Notably, by allowing trait-mediated biotic interactions to directly influence biogeographic evolution, we are able to recover evidence for competitive exclusion during the radiation of Darwin’s Finches (Figure 6). That is, niche dissimilarity facilitated the colonization of new areas during the finch radiation. We observe that all four traits shaped the rates of colonization, to different extents, among the different islands in the archipelago. This is in accordance to theoretical and other empirical evidence suggesting that coexistence can only be tenable with some degree of niche divergence (Elton 1946; Hardin 1960; Macarthur and Levins 1967; Diamond 1978; Godoy et al. 2014). Furthermore, since successful colonization is a necessary step to increase an area’s biodiversity, our results hint at the mechanism in which microevolutionary processes might lead up to macroevolutionary patterns, such as the generation of spatial variation in richness. More detailed inspection of per lineage-area effects of biotic interactions during the clade’s evolutionary history allows us to disentangle between biogeographic events that involved biotic interactions against those that did not (Figure 6).

### Inferring trait-range histories

The development of phylogenetic models has allowed researchers to reconstruct historical processes, even when restricted to only extant information, and to test central hypotheses regarding the tempo and mode of evolutionary dynamics (Garamszegi 2014). Such models are valuable, in part, because they require hypotheses about the mode by which lineages evolve and diversify (e.g., Butler and King 2004) to be defined in formal terms (e.g., in an SDE). Understanding what features are and are not formally modeled determines what one may prudently conclude from analyses under the method, which we aim to make explicit below. While our model entails several simplifying assumptions, future work may relax these assumptions to incorporate additional features important to modeling trait and range co-evolution.

The simple biogeographic model used here assumes that at the moment of speciation the daughter lineages inherit identical ranges as their ancestor lineage, a particular case of sympatric speciation. Given that the great majority of speciation events involve a phase of geographical isolation (Mayr 1970; Rundell and Price 2009), we acknowledge that this assumption does not hold in most empirical systems. Importantly, by not allowing allopatric cladogenesis *sensu* Ree et al. (2005), the inferred parameters governing biotic interactions can be equivocal on a clade with a history of allopatric speciation. For instance, the effect of competitive exclusion (*ω*_1_) is presumed to be large between recently diverged species, yet, these are forced to coexist instantly after speciation, probably underestimating the effect of similarity in colonization rates (e.g., secondary contact times) by overestimating the period of sympatry and bearing upon *in situ* biotic interactions (*ω*_*x*_) to explain the trait variance. Consequently, an important next step is to incorporate models that allow for different modes of geographical speciation, such as the Dispersal-Extinction-Cladogenesis (DEC) model and relatives (Ree et al. 2008; Matzke 2014). This requires designing efficient data augmentation proposals and their associated Metropolis-Hastings ratios, which we are currently working to solve. Other relevant biogeographic processes are not being considered, but are relatively straightforward to incorporate in future versions of the model. Characteristics of the delimited geographical regions, such as distance from each other (Landis et al. 2013), geographical area (Tagliacollo et al. 2015), connectivity (Kadmon and Allouche 2007), age of area availability (e.g., on volcanic islands, Landis et al. 2018), and resource availability (Tilman 1985) will provide key information when inferring biotic interactions. Furthermore, incorporating abiotic optima, as determined by the different regional environments, could be used to distinguish abiotic from biotic forces acting upon trait and range evolution. Research in these directions would further demonstrate the potential of inferring trait and biogeographic evolution as interacting processes (Sukumaran and Knowles 2018).

Assuming that interspecific competition acts upon only a single axis of niche evolution, as we assume, may be problematic (Connell 1980). Species niches are better thought of as multidimensional hypervolumes (Hutchinson 1957), and so viewing this complexity through a single, univariate trait must misrepresent the true nature of biotic interactions between species (Diamond 1978; Grether et al. 2009). In some cases, fitting the model separately to each trait or asserting independence on the traits by multivariate transformations (such as PCA) can unduly influence parameter estimates (Uyeda et al. 2015; Cadena et al. 2018). For example, a lack of evidence for biotic interactions within a given axis does not rule out competition from occurring along other unmeasured resource utilization axes (Connell 1980). We advise the researcher to select a trait of study that has been suggested as relevant to niche partitioning (e.g., bill size and shape in the Darwin’s finches; Grant and Grant 2002). Measuring species niche overlap between partitions, however, is a general problem pervasive across ecology (Diamond 1978; Petraitis 1979).

Species usually occupy ranges of values along niche axes (e.g., the range of temperature where the species can persist) or have considerable intraspecific variation; these features warrant modeling in future methods (e.g., as in Quintero et al. 2015). Moreover, niche similarity might differ between univariate and multivariate spaces, and improved phylogenetic models of competition should account for the multivariate distances between value ranges in niches (Huelsenbeck and Rannala 2003). Despite complications in identifying and representing which traits may be involved in competition, competitive forces are thought to be stronger among recently diverged species because of their overall similarity in resource use (Darwin 1859). Likewise, we assume that biotic interactions have had the same directionality and magnitude (relative to phenotypic dissimilarity) across all lineages throughout the clade’s evolutionary history, even though the magnitude and sign of competitive effects probably varies within and between clades, contingent on measured, unmeasured, and unknown factors. While our current model tests for the constant effect of a clade-wide competitive process influencing a univariate trait, it may be extended to accommodate multivariate traits, trait value ranges, and branch-heterogeneous competitive effects.

Finally, our model assumes that biotic interactions only occur between lineages modeled by the phylogenetic tree, which we take to be the reconstructed tree—a tree that only represents lineages corresponding to the set of most recent common ancestors shared among the sampled taxa. Modeling competition while naively taking the reconstructed tree to represent the true evolutionary history among all lineages overlooks any historical contribution from lineages left absent in the reconstructed tree, namely absent lineages representing the ancestors of excluded, unsequenced, or extinct lineages. While, in principle, we can improve representation among extant lineages, that is not always the case with extinct lineages, yet disregarding the influence of extinct lineages is known to mislead some evolutionary inferences (Schindel and Gould 1977; Slater et al. 2012). Being blind to paleobiological interactions may be particularly troublesome in our case, since the geographic and phenotypic evolution of any one ancestral lineage should depend on that of all other contemporaneous lineages, independent on their survival to the present. Provided the data are available, spatial and morphological information from paleontology could be incorporated in our model to attain more biological realism and broaden applicability to clades were extinction rates have been presumed to be high (Mitchell 2015). Correctly modeling the influence of competitive effects with extinct or unsampled ghost lineages that are not represented in the model will require the the introduction of features from birth-death processes.

At first glance, developing such a model appears mathematically and methodologically challenging, but progress here would be rewarding. Modeling interactions between trait evolution, competition, biogeography, and diversification processes in a phylogenetic context would represent a major advance towards how we understand the generation and maintenance of biodiversity. As phylogenetic models of competition continue to mature, we must strive to incorporate trait-diversification dynamics that are thought to underlie well-studied macroevolutionary phenomena, such as the Great American Biotic Interchange (GABI; Simpson 1950; Benton 1987). The biogeographic exchange of lineages during GABI is considered to be the result of competition between distantly related clades (Diamond 1978), and classic macroevolutionary hypotheses, such as the “Red Queen” (Van Valen 1973), suggest that temporal and spatial turnover in taxa results mostly from biotic interactions.

### Bayesian data augmentation

In our work, we provide a framework to test the effect of ecological processes on phenotypic and biogeographical distribution of species across evolutionary time. The Bayesian data augmentation framework we present here is robust yet flexible, making it adaptable to similar inference problems of associated discrete and continuous character co-evolution. For instance, similar models were developed for processes of correlated nucleotide substitution rates and Brownian motion evolution (Lartillot and Poujol 2011; Horvilleur and Lartillot 2014; Lartillot et al. 2016), and it is conceivable that nucleotide substitution patterns should in some way reciprocally influence how molecular phenotypic traits, such as protein function, evolves (Robinson et al. 2003; Rodrigue et al. 2006). We hope that our algorithmic framework encourages and allows other researchers to develop phylogenetic models that study the interdependent effects of continuous and discrete trait evolution within and between lineages.

## Conclusion

Given the ubiquity of character displacement, it might be tempting to assume that phenotypic divergence is the direct result of natural selection acting to avoid competition on sympatric populations (Grant 1972). But it is also plausible that those populations were only able to spread into sympatry because their niche was sufficiently different in the first place (Schluter and McPhail 1992). Lack (1954) pointedly outlined this difference over half a century ago when discussing a case of the bird genus *Sitta*: “… the two species show no overlap in beak measurements [where they occur in sympatry], a difference presumably evolved through the need for avoiding competition for food; or rather, it is only where such a difference has been evolved that the two forms can live alongside each other”. Jointly examining distinct mechanisms in trait and biogeographic evolution allows testing core evolutionary theories on how biodiversity is brought about. Clearly, the process by which species diversify phenotypically and attain coexistence is fundamentally important to the generation of spatial gradients of diversity, and thus further understanding the underlying mechanisms is a paramount goal of evolutionary biology.

## Acknowledgments

We are grateful for conceptual feedback provided by Michael Donoghue that helped improve the clarity and presentation of the manuscript, to Nathan S. Upham who provided feedback on earlier versions of the work, and to Mike R. May for his thoughts on how to improve the description of methods.

